# The diversification of termites: inferences from a complete species-level phylogeny

**DOI:** 10.1101/2020.12.07.414342

**Authors:** Marcio R. Pie, Tiago F. Carrijo, Fernanda S. Caron

**Affiliations:** Departamento de Zoologia, Universidade Federal do Paraná, C.P 19020, Curitiba PR 81531-990, Brazil; Centro de Ciências Naturais e Humanas, Universidade Federal do ABC, São Bernardo do Campo, Brazil

**Keywords:** Isoptera, Termitidae, diversification, speciation, TACT, phylogenetic imputation

## Abstract

Termites play a major role in a variety of ecological processes in tropical and subtropical biomes worldwide, such as decomposition, soil formation and aeration, and nutrient cycling. These important ecosystem services were achieved through their highly complex societies and remarkable adaptations, including the evolution of sterile worker castes, the acquisition of endosymbionts, and the capacity for extensive environmental engineering, yet the causes and consequences of their ecological success are still poorly understood. The goals of our study were (1) to provide the first complete, species-level phylogeny of all currently recognized termite species by integrating the available genetic and taxonomic data, as well as methods of phylogenetic imputation and divergence time estimation; and (2) to explore variation in speciation rates among termite lineages. We provide the inferred relationships as a set of 1,000 pseudo-posterior trees, which can be used in future comparative analyses. We demonstrate that speciation rates have been relatively constant throughout the history of termites, with two positive shifts in speciation rates: one at their origin of Euisoptera and the other concordant with evolution of Termitidae. On the other hand, there was no obvious trend towards deceleration in speciation rates for termites as a whole, nor within the most species-rich families. The provided trees might represent a valuable resource for termite comparative studies by summarizing the available phylogenetic information, while accounting for uncertainty in the inferred topologies.

## 1. Introduction

Termites are the quintessential environmental engineers in the tropics, performing many important ecosystem services such as decomposition, soil formation and aeration, and nutrient cycling (Jouquet et al., 2006, 2011; Evans et al., 2011; Ashton et al., 2019). They are dominant animals in the soil, comprising up to 40–65% of the total soil macrofaunal biomass (Eggleton et al., 1996; Tuma et al., 2020), and can be responsible for 54–68% of total decomposition in some biotopes (Jouquet et al., 2011; Ashton et al., 2019). Despite the important ecological role of termites on a variety of terrestrial ecosystems, our understanding of the adaptations that led to such high prevalence is still limited (Davis et al., 2009; Legendre and Condamine, 2018). Investigating the causes and consequences of the remarkable ecological success of termites will require not only extensive ecological and life-history data, but also a broad understanding of their phylogenetic relationships.

The evolutionary route leading to eusociality in termites was different from that of other eusocial insects, with ancestral termites probably evolving from subsocial organisms similar to present-day *Cryptocercus* cockroaches (Cleveland et al., 1934; Legendre and Grandcolas, 2018). Relatively little is known about termite diversification other than inferences on the timing of the most basal splits (e.g. Ware et al., 2010; Bourguignon et al., 2015, 2017; Legendre et al., 2015). To the best of our knowledge, only three studies to date used analytical methods to quantify patterns of termite diversification. Davis et al. (2009) tested the conjecture postulated by Wilson (1992) that the evolution of social behavior would be associated with a decrease in diversification rates. No evidence for such a decrease was found by Davis et al. (2009), and a heterogeneity in diversification rates among termite lineages was revealed, yet the low power provided by the use of sister-clade comparisons necessarily limits the scope of their analyses. More recently, Legendre and Condamine (2018) used parametric methods to explore large-scale diversification patterns in Dictyoptera and found evidence for two main rate shifts: one at the origin of termites (with eusociality) and another at the origin of Termitidae (with the loss of cellulolytic parabasalids symbionts). They also found that lineages with true workers diversified faster than those with pseudergates (“false” workers). True workers are individuals that diverge early in the developmental pathway from the imaginal line, while pseudergates have high developmental plasticity, being able to undergo progressive, stationary, and regressive molts, eventually becoming alates (Korb and Hartfelder, 2008). This trait is often related to the degree of eusociality in termites, and species with true workers are considered more highly eusocial (Shellman-Reeve, 1997). Finally, Condamine et al. (2020) used an extensive analysis of fossil and phylogenetic data to explore temporal patterns of diversification in Dictyoptera, particularly with respect to the potential impacts of mass extinctions. Although their results are comprehensive and robust, they tended to focus at the level of Dictyoptera and, given that it includes groups that are vastly different in their biology, it is not clear to what extent they would reflect specifically the trends within Isoptera. Nevertheless, despite these important contributions, more fine-scale variation in diversification among termite lineages is still poorly understood.

Important advances have been obtained for termite systematics in recent years using morphological and mitogenomic data (Inward et al., 2007; Ware et al., 2010; Cameron et al., 2012; Bourguignon et al., 2015, 2017; see also Legendre et al., 2015). More recently, Bucek et al. (2019) inferred the main relationships among 66 termite lineages using transcriptome data, and the results were largely congruent with previous hypotheses. However, the availability of phylogenetic information in termites (as in most taxa) is highly uneven, such that relatively few lineages account for most of the sequences available in public repositories, whereas little or no information is available for many species. Given that many types of inference require the use of large-scale phylogenies (such as diversification studies, see Pie and Tschá, 2009; Chang et al., 2020), one important alternative is the integration of phylogenetic and taxonomic data with simple models of lineage diversification to provide pseudo-posterior trees that can be used in downstream analyses. In particular, analyses can be repeated with many alternative topologies, thus accounting for phylogenetic uncertainty and therefore allowing for inferences that would not be feasible based on taxonomically-limited trees (e.g. Arnan et al., 2018). In this study we integrate phylogenetic and taxonomic data to provide the first-species level phylogeny of all currently recognized termite species. In addition, we use the inferred relationships to explore variation in speciation rates among termite lineages.

## 2. Methods

We obtained complete trees for all currently valid termite species using a two-stage approach that takes as inputs backbone topologies based on molecular data and sets of taxonomic constraints (e.g. placing species within their corresponding genera, subfamilies, or families) using Taxonomic Addition for Complete Trees: TACT (Chang et al., 2020). TACT is similar to another two-stage approach known as PASTIS (Thomas et al., 2013), which was used in previous studies to provide large-scale trees of several taxa, including birds (Jetz et al., 2012), squamates (Tonini et al., 2016), and sigmodontine rodents (Maestri et al., 2017). The main advantage of TACT in relation to PASTIS is that it provides branch lengths based on local diversification rates, as opposed to global rates, thus being more suitable for large phylogenies with heterogeneous rate regimes (Rabosky et al., 2018; Chang et al., 2020; Siqueira et al., 2020). Thomas et al. (2013) categorized species into three types: type 1 species have genetic information; type 2 species have no genetic information but are congeners of a species with genetic information; and type 3 species have no genetic data and are members of a genus that does not have genetic data. In our dataset we have 402, 2017, and 550 species from each type, respectively. The main steps in our phylogenetic analyses are shown in Figure 1.

**Figure 1.**
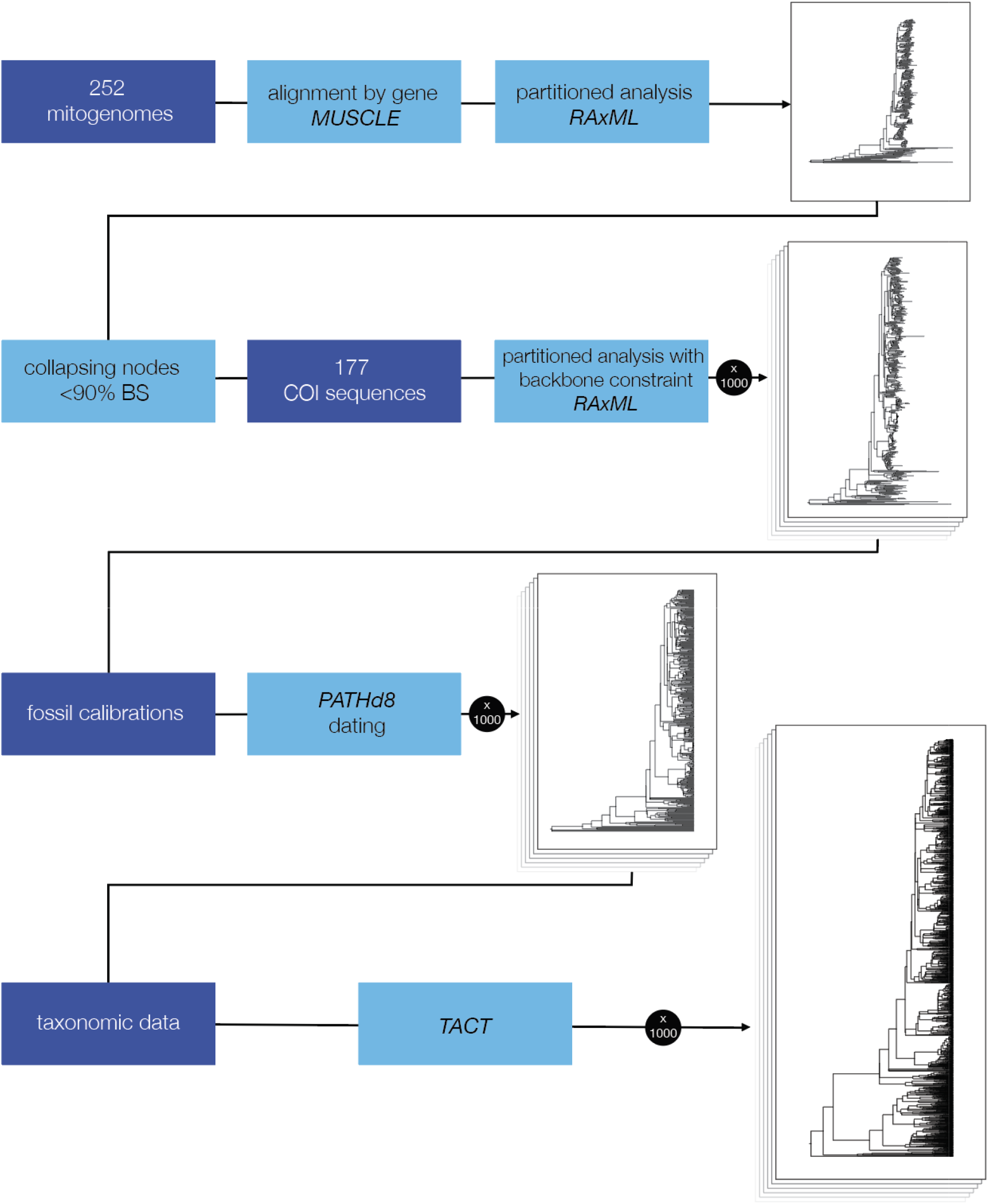
Diagram indicating the main steps in our analyses. Dark blue boxes indicate sources of data, whereas light blue boxes correspond to analyses. TACT: Taxonomic Addition for Complete Trees.

Our backbone trees were obtained by combining information on 252 mitogenomes and 177 COI sequences available on GenBank and BOL (Table S1). Although other loci are available for a reduced number of termite species, we chose to focus only on COI and complete mitogenomes to avoid the computational challenges involved in sparse matrices for this level of taxon sampling. Likewise, we chose not to include the transcriptome data from Bucek et al. (2019) given that their phylogeny is largely congruent with previous mitogenomic studies. The only exception is the placement of Sphaerotermitinae as sister clade to Macrotermitinae, which was not uncovered with confidence in previous studies. We therefore used this relationship as a topological constraint in our phylogenetic analyses. Each locus was aligned separately using MUSCLE (Edgar, 2004) and then concatenated. The best model of evolution for the dataset was determined using PartitionFinder 2 (Lanfear et al., 2016). Preliminary analyses using the complete dataset showed erratic behavior because of some species for which there was only COI data. Therefore, to improve the quality of the backbone trees, we first ran a partitioned analysis on RAxML using only mitogenomic data and retained only the nodes with over 90% bootstrap support (Figure S1). We then enforced the monophyly of the genera on the mitogenomic + COI dataset, as well as the placement of Sphaerotermitinae indicated above, and repeated the partitioned RAxML with the full dataset, retaining 1000 bootstrap trees for downstream analyses (Figure S2). We used the fossil calibrations indicated in Tables S2-S5 to obtain calibrated trees using PATHd8 v. 1.0 (Britton et al., 2007). Minimum and maximum age constraints were specified for all calibrations, except for the oldest, as PATHd8 requires a fixed age, and the average between the minimum and maximum age constraints was used. PATHd8 is a rate-smoothing method similar to non-parametric rate smoothing (Sanderson, 1997) that allows for the efficient estimation of divergence times for large trees. We decided to use this combination of RAxML and PATHd8 because preliminary analyses using BEAST 2.6.1 (Bouckaert et al., 2019) showed considerable difficulties to obtain convergence on this dataset. The divergences in the resulting trees was summarized using TreeAnnotator v.2.6.3 (Rambaut & Drummond 2020) using the settings to generate a maximum clade credibility tree with median node heights. There were a few isolated cases of negative branch lengths, possibly due to high variance in estimates of divergence times in particular nodes (see Heled & Bouckaert 2013). Instead of enforcing all branch lengths to be positive, we left those branches in the summary tree to allow for the reader to identify their location and interpret them with the corresponding caution. However, it is important to note that this phenomenon is only present in the summary tree and not in the individual trees used as input.

Taxonomic data for all currently recognized termite species were obtained using the Termite Database (Constantino, 2016; last accessed on June 10, 2020). We included as taxonomic levels their corresponding genera, families and (in the case of Termitidae and some Rhinotermitidae), subfamilies. To some well-established non-monophyletic taxa, such as Rhinotermitidae, Termitinae and *Nasutitermes*, no constraint was set. Species were added to each of the 1000 backbone trees indicated above to generate 1000 pseudo-posterior trees. It is important to note that nine of the mitogenomes were not identified to species; they were still used in TACT to assist in the placement of species in their corresponding lineages, but were pruned afterwards.

We explored variation in speciation rates among termite lineages using the DR statistic (Jetz et al., 2012; hereafter, λDR), as implemented in *ES-sim* (available at https://github.com/mgharvey/ES-sim). λDR is a non-model-based estimator of speciation rate that is calculated as a weighted average of the inverse branch lengths between a given species to the root of the phylogeny (i.e. the root-to-tip set of branches). It is therefore similar to the node-density estimator (Freckleton et al., 2008), except that it places more emphasis on recent branch lengths (Jetz et al., 2012). As a consequence, λDR tends to reflect speciation rates rather than net diversification rates (Jetz et al., 2012; Belmaker and Jetz, 2015). We calculated λDR for each of 500 pseudo-posterior trees and visualized variation in λDR among lineages using the contMap function in *phytools* 0.7-47 (Revell, 2012) in R 4.0.1 (R Core Team, 2020). We also explored diversification patterns in deeper nodes of the tree using lineages-through-time (LTT) plots of all termites, as well as specific clades (Termitidae, Kalotermitidae, Apicotermitinae, Macrotermitinae, Nasutitermitinae, Cubitermitinae). We carried out LTTs with three different trees: (a) trees including only species with genetic data; (b) trees augmented using TACT but that were all based on the same MCC guide tree, and (c) the trees in the first group that were augmented using TACT. These three LTTs therefore allow for assessing uncertainty due to phylogenetic inference, taxonomic augmentation, and both factors, respectively.

## 3. Results

A summary tree of the inferred relationships and divergence times for species with genetic data, after calibration using PATHd8, is shown in Figure 2. According to these estimates, the most common recent ancestor of all termites evolved during the early Jurassic Period, whereas Neoisoptera arose between the early and late Cretaceous (Figure 2). Sensitivity analyses using four alternative calibration schemes (Table S2-S5) provided very similar results (Figure 2A).

**Figure 2.**
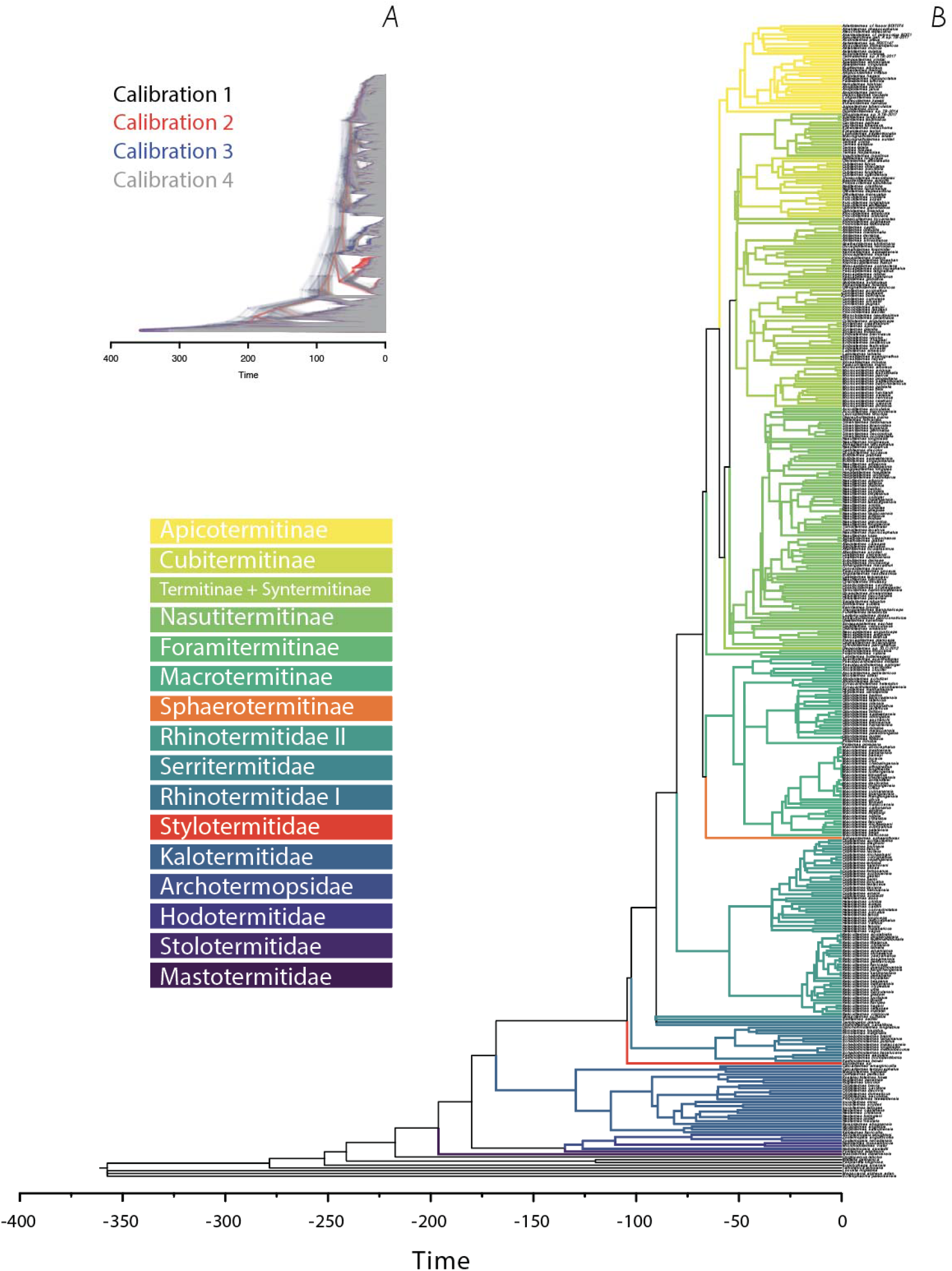
Divergence time estimates of termite lineages with available genetic data. A. Densitree of four different sets of trees indicating how the resulting estimates were largely congruent between calibration schemes (Table S2-S5); B. Summary tree of 1000 constrained bootstrap trees after being ultrametricized using PATHd8 using calibration 3 (Table S4). See text for details.

The mapping of speciation rates (as measured by the λDR statistic) onto the termite tree was able to reveal the main diversification regimes during the history of termites (Figure 3). The families Mastotermitidae, Archotermopsidae, Hodotermitidae, Stolotermitidae and Kalotermitidae in general were characterized by low speciation rates. A few lineages showed an increased diversification rate within these clades, and these patterns are related to the origin of some genera, such as *Anacanthotermes* (Hodotermitidae) and *Bifiditermes* (Kalotermitidae). Within the non-Termitidae Neoisoptera, the groups Serritermitidae, Rhinotermitinae, *Termitogeton* and some clades of *Heterotermes* also showed lower speciation rates. Specially Serritermitidae, which the evolutionary regime is comparable only with that of Mastotermitidae and Archotermopsidae. Stylotermitidae and *Coptotermes* had similar regimes, and *Reticulitermes* had a fast radiation in the last 23 Mya.

**Figure 3.**
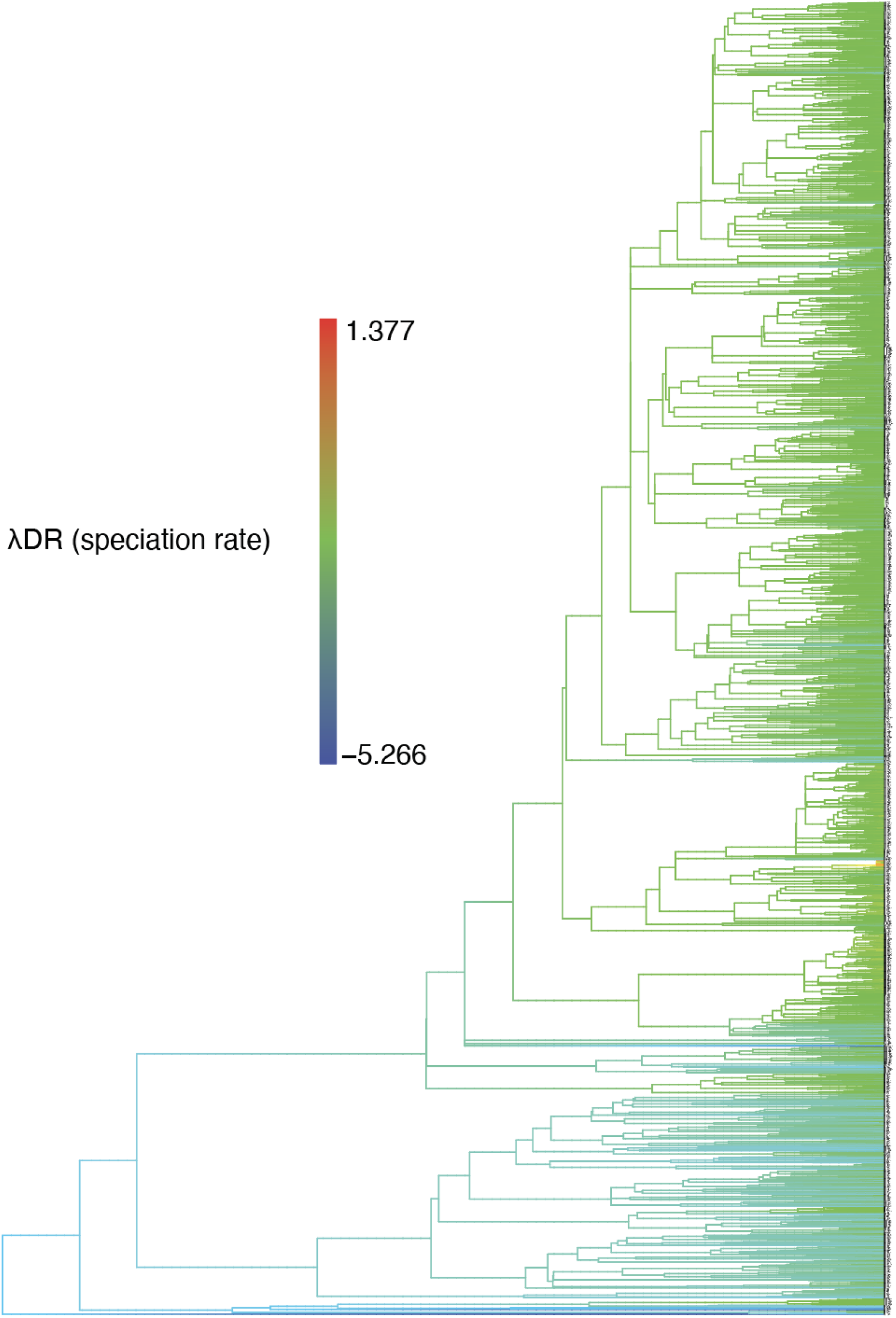
Phylogenetic relationships among all currently-recognized termite species. Branches were colored based on their corresponding speciation rates.

As expected, most Termitidae clades showed relatively high speciation rates, but a few different evolutionary regimes can be observed among lineages. The fungus-growing termites (Macrotermitinae) had the fastest radiation considering all termites. This subfamily originated ca 70 Mya, but the evolutionary regime speeded up in the last 10-20 Mya, especially within the genus *Macrotermes* (Figs. 3, 4). Nasutitermitinae also showed an increase in speciation rate in the last 10 Mya, that can be observed through the LTTs plots (Fig. 4). Apicotermitinae and Cubitermitinae, on the contrary, had relatively constant rates since their origin, and the Foraminitermitinae clade showed the lowest speciation rate within Termitidae.

**Figure 4.**
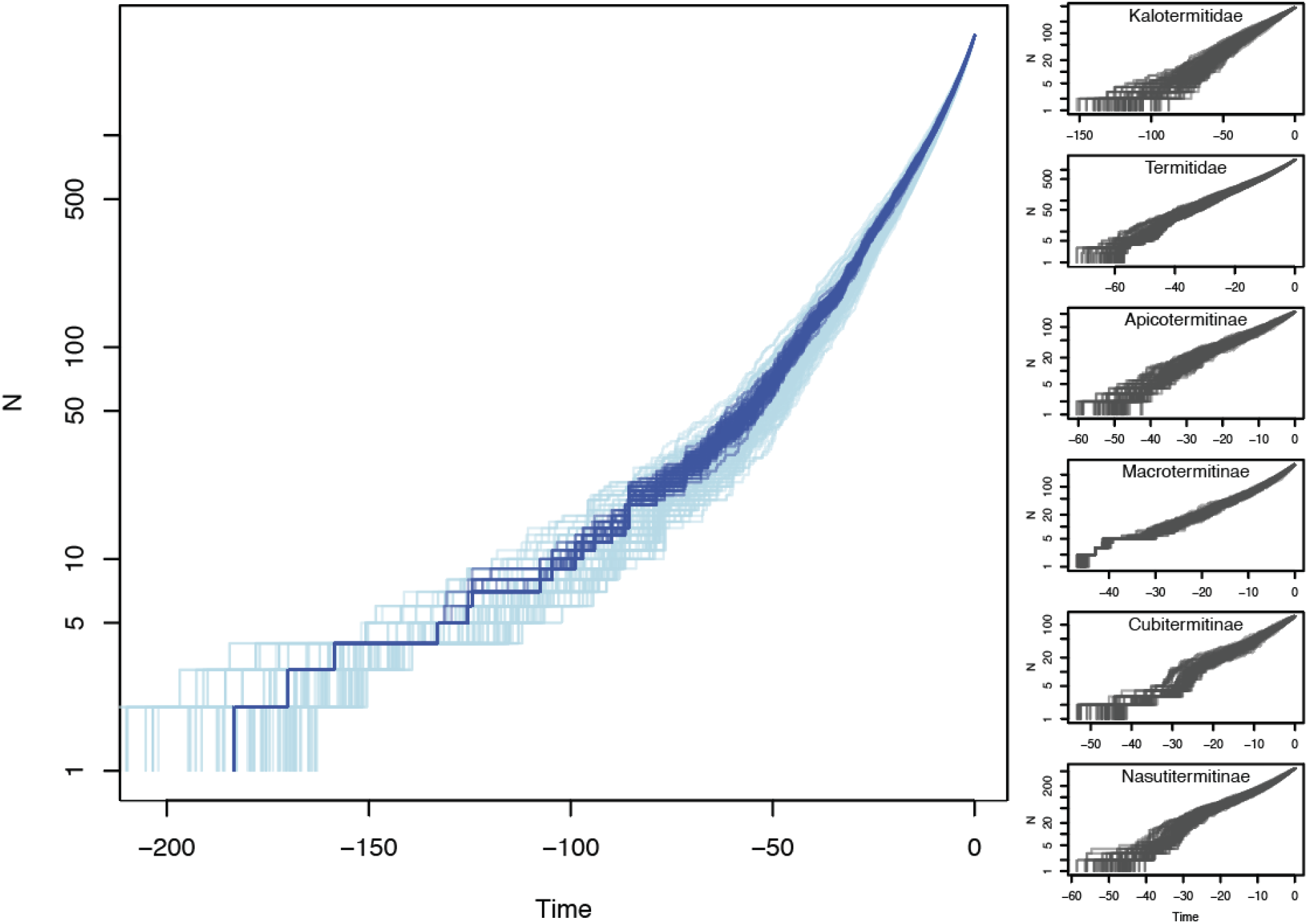
Lineages-through-time (LTTs) plots of 100 pseudo-posterior topologies of all termites (dark blue), as well as 100 pseudo-posterior topologies based on TACT imputation on the same backbone topology (light blue). Figures on the right correspond to LTTs of specific clades, namely Kalotermitidae, Termitidae, Apicotermitinae, Macrotermitinae, Cubitermitinae, and Nasutitermitinae.

## 4. Discussion

In general, our results with respect to the main relationships are particularly congruent with those obtained by Bourguignon et al. (2015, 2017), but there was broad agreement between our results and those of several published studies: (1) *Mastotermes* as sister to all other termites (Inward et al., 2007; Ware et al., 2010; Cameron et al., 2012; Bourguignon et al., 2015; Legendre et al., 2015; Bucek et al., 2019); (2) Archotermopsidae as paraphyletic with respect to Hodotermitidae (Legendre et al., 2015); (3) Kalotermitidae as sister to Neoisoptera (Ware et al., 2010; Legendre et al., 2015; Bucek et al., 2019); (4) Rhinotermitidae as paraphyletic, with respect to Serritermitidae and Termitidae (Inward et al., 2007; Legendre et al., 2015; Bourguignon et al., 2015); (5) Heterotermitinae (including Coptotermitinae) as sister group to Termitidae (Bourguignon et al., 2015; Bucek et al., 2019); (6) Macrotermitinae and Apicotermitinae monophyletic (Bourguignon et al., 2017; Inward et al., 2007; Legendre et al., 2015); (7) Termitinae paraphyletic with respect to Syntermitinae, Nasutitermitinae and Cubitermitinae (Bourguignon et al., 2017; Bucek et al., 2019; Legendre et al., 2015).

Our time-calibrated tree estimated termites to have originated between 176 and 232 Mya and Termitidae between 60 and 80 Mya. While these dates are congruent with Ware et al. (2010), most recent works have estimated more recent origins to these clades, roughly 50 Mya more recent to Isoptera and 20 Mya to Termitidae (Engel et al., 2009; Legendre et al., 2015; Bourguignon et al., 2015, 2017; Bucek et al., 2019). The causes of such discrepancy are still unclear. Further exploration of divergence time estimates, particularly using more powerful methods such as the fossilized birth-death approach (Heath et. 2014) and that include more unlinked nuclear loci might be particularly revealing.

*Mastotermes darwiniensis*, the sole living species of Mastotermitidae and an ancient lineage with a single living species from Australia, showed a very low diversification rate. This agrees with the diversification shift found by Davis et al. (2009) between Mastotermitidae and the other termite families. However, the fossil record of Mastotermitidae is rich and distributed all over the world (Zhao et al., 2019; Bezerra et al., 2020), and this pattern may be for a high extinction rate that followed a Cretaceus radiation of the group (Engel et al., 2009). Within the clade composed by Hodotermitidae, Stolotermitidae and Archotermopsidae, the genus *Anacanthotermes* shows a singular evolutionary regime, with a high speciation rate in the last 25 Mya. Along with the other two living genera of Hodotermitidae, these are harvest termites, and our results are in agreement with the fossil record of the family, suggesting that this clade originated and expanded with the grasslands during the Oligocene and Miocene (Krishna et al., 2013).

Both *Mastotermes* and Hodotermitidae are considered to have a true worker caste (Korb and Hartfelder, 2008), and Legendre and Codamine (2018) found that societies with this innovation showed a high overall diversification rate. As discussed above, however, these two groups showed different evolutionary regimes in our analysis, and the true worker alone is probably not a sufficient explanation for the diversification patterns. In fact, the definition of true workers is often associated with two independent traits: (1) sterility, and (2) developmental constraint to the imaginal line. In *Mastotermes*, workers have a developmental constraint, but are not sterile (Bourguignon et al., 2016), whereas in Hodotermitidae, as well as in most of the termites that showed a high speciation rate (see below), a (true) sterile worker caste is present. Thus, worker sterility alone may be a better explanation for termite evolutionary patterns, once a highly altruistic worker caste is a good indicator of the spectrum of eusociality in these insects (Shellman-Reeve, 1997).

The family Kalotermitidae is the second largest among termites in terms of number of species, and such richness alone probably led Davis et al. (2009) to recover in their analysis a diversification shift in the origin of the clade Kalotermitidae + Neoisoptera. Our results, however, showed a low overall speciation rate for this family (Fig. 2). In comparison to other groups with similar numbers of living species, such as Nasutitermitinae or Macrotermitinae, this is an old clade with a constant speciation rate over time (Fig. 3), and the currently high diversity may represent low extinction rates instead of a high speciation rate. Also, colonization by rafting is favored by the wood nest habits of Kalotermitidae, and many species of this group are important components of oceanic islands faunas. These dispersion/colonization events were probably important drivers of diversification for the group (Krishna et al., 2013).

The genera *Reticulitermes* and *Stylotermes* showed evolutionary regimes with high speciation rates, but both may have been overestimated by a taxonomic bias. *Stylotermes* and *Reticulitermes* comprise 45 and 142 described species, respectively, but the validity of many of these names are questionable, and many synonyms may occur after taxonomic revisions (Takematsu and Vongkaluang, 2012; Wu et al., 2018). Taking in consideration this bias, the main speciation shift within termite evolution was the origin of Termitidae (Figs. 2, 3; see also Davies et al., 2009; Engel et al., 2009; Legendre and Codamine 2018). The radiation of this clade is related to the loss of the symbiotic relationship with the cellulolytic parabasalids, those same that were one of the main factors driving the origin and radiation of earlier termites (Higashi and Abe, 1997; Nalepa, 2015). Bucek et al. (2019) proposed that feeding on this organic rich substrate starved the cellulolytic parabasalids symbionts to extinction. This ended up allowing them to have a diverse diet, including the entire decomposition gradient of plant material, from sound wood, to grass, epiphytes, to soil and cultivation of fungi simbiontes (see also Engel et al., 2019). This diversification made the Termitidae family to account for the impressive biomass of termites in the tropics and subtropics, compasing almost 80% of all described termite species (Engel et al., 2009; Tuma et al., 2020). Also, nearly half of the termite species are soil feeders, and this habit was crucial to the radiation of the family (Higashi and Abe, 1997; Krishna et al., 2013; Bucek et al., 2019; Engel et al. 2019).

Macrotermitinae was the clade with the highest speciation rate in our results, and the external rumen with cultivation of *Termitomyces* fungi (basidiomycetes) was probably the driving force to the ecological success of this subfamily in the Old World. Within this clade, yet two patterns could be observed (Figs. 2, 3): a first one at the origin of the clade, and the second in the last 10 Mya, probably related to the savannas and C4 grasse expansions in Africa (Edwards et al., 2010; Nobre and Aanen, 2012; Charles-Dominique et al., 2016).

The Nasutitermitinae subfamily originated 57 Mya and showed a high speciation rate throughout the time (Fig. 2, 3). Specialization of a strict chemical defense in this clade is often regarded to evolve driven by an army-race against termite’s major predators, the ants (Higashi and Abe, 1997; Engel et al.m 2009). The increase in the fossil record of ants in the last 50-60 Mya and our results for the origin of this group is consistent with this hypothesis (Ward, 2004).

It is important to emphasize that the stochastic polytomy resolution approach used in this study has limitations. First, the level of phylogenetic information necessarily varies across the tree, with some clades having more or less underlying data. This effect is nearly unavoidable, given that the choice of taxa for phylogenetic studies is never random—researchers tend to include phylogenetically distant species and/or species that are available for sequencing. However, the pattern of accumulation of lineages (Figure 3) seems fairly robust despite the uncertainty involved in phylogenetic imputation. Nevertheless, we expect that this limitation might become increasingly less severe as additional species are sequenced and included in future studies. Another important limitation is that the analysis is necessarily constrained to include species that have been described. The extent to which different termite clades are more or less susceptible to this bias is currently unknown and possibly unknowable. Despite these limitations, we envision the use of large-scale phylogenetic information provided in our study might serve as an important tool to uncover a variety of phenomena relevant to termite ecology and evolution.

## Acknowledgments

This study was partially funded through a grant from CNPq (#301636/2016-8) to MRP.

## Supplementary material

**Table S1**. Accession numbers of the sequences obtained from the GenBank or IBOL databases and used in the present study.

**Table S2**. Calibration scheme used in the present study as calibration 1.

**Table S3**. Calibration scheme used in the present study as calibration 2.

**Table S4**. Calibration scheme used in the present study as calibration 3.

**Table S5**. Calibration scheme used in the present study as calibration 4.

**Figure S1**. Relationships between termite lineages based on complete mitogenomic data. Node values correspond to bootstrap support. Branches with bootstrap support <90% were collapsed.

**Figure S2**. Relationships between termite lineages based on COI sequence data and constrained by the topology indicated in Figure S1. Node values correspond to bootstrap support.

