## Supplementary figures and images for "The diversification of termites: inferences from a complete species-level phylogeny"

### Figure S1

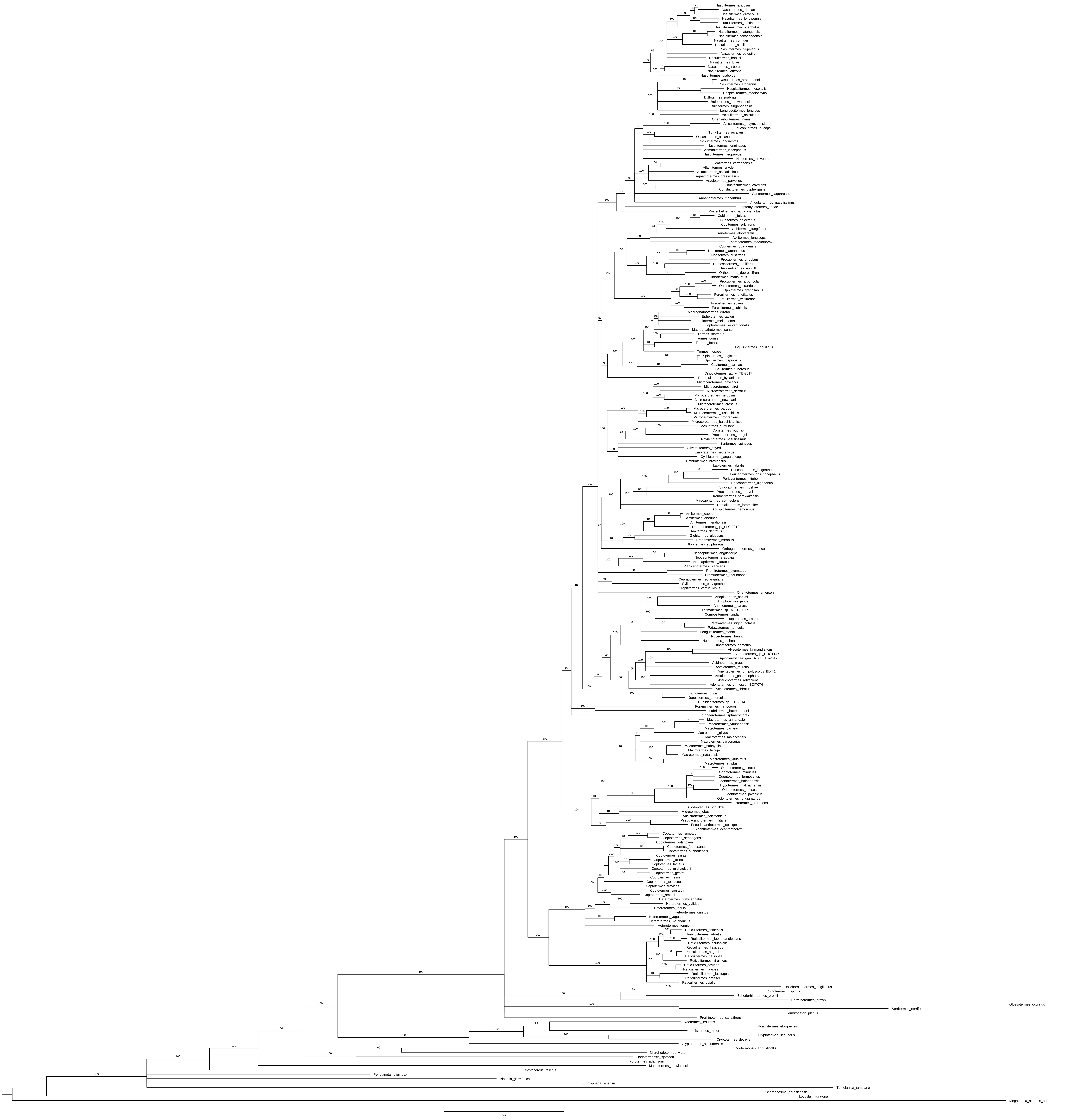

### Figure S2

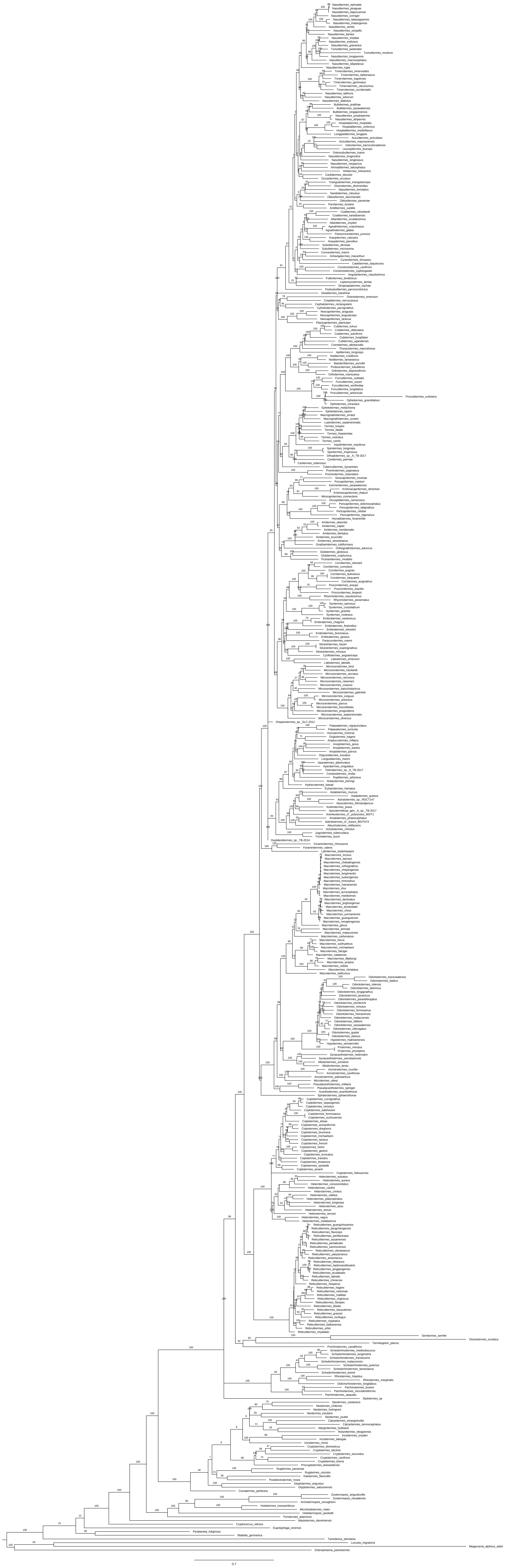
